# Warming sensitivity of spring phenology of deciduous species lies in their bud energy budget

**DOI:** 10.1101/2021.05.03.442458

**Authors:** Marc Peaucelle, Josep Peñuelas, Hans Verbeeck

**Affiliations:** Computational and Applied Vegetation Ecology - CAVElab, Department of Environment, Faculty of Bioscience Engineering, Ghent University, Gent, Belgium; CSIC, Global Ecology Unit CREAF-CSIC-UAB, 08193 Bellaterra, Catalonia, Spain; CREAF, 08193 Cerdanyola del Vallès, Catalonia, Spain

**Keywords:** plant phenology, budburst, temperature, light, energy budget, modelling, climate warming

## Abstract

Spring phenology is mainly driven by temperature in extratropical ecosystems. Contrasting responses of foliar phenology to climatic warming, however, have been reported in recent decades, raising important questions about the role of other environmental constraints, especially light. In fact, temperatures differ substantially between plant tissues and the air because plants absorb and lose energy. Yet, phenology studies always substitute plant tissue temperature by air temperature. Here, we explored how solar radiation, wind, and bud traits might affect spring phenology of deciduous forests through the energy budget of buds. We show that air temperature might be an imprecise and biased predictor of bud temperature. Our current interpretation of the plant phenological response to warming should be reconsidered, which will require new observations of bud traits and temperature for accurately quantifying their energy budget.

---

Plant phenology, the study of the timing of life-cycle events, drives several ecosystem functions, such as plant productivity and biomass, but also local and global climates by affecting biogeochemical and biogeophysical processes, such as carbon storage and energy fluxes^1,2^, and the abundance and diversity of local flora and fauna, such as pollinators and herbivores^3,4^. Understanding the environmental controls and responses of plant phenology to climate change is thus essential for several sectors, e.g. agriculture, forestry and gardening^5^, but also for conservation^6^ and public health^7^ (e.g. allergies).

## Foliar phenology and temperature

Leaves control plant water loss and carbon assimilation and are thus central to the growth of other plant organs. Temperature is one of the main drivers of foliar phenology in extratropical ecosystems^8^. Cold temperatures during winter and warm temperatures during spring control leaf unfolding and flowering at the beginning of the season. Temperature during the growing season will control foliar development, carbon assimilation, transpiration, plant growth and the establishment of new buds for the following year. Finally, cold temperatures at the end of the growing season are sensed by plants as a signal for foliar senescence^9^.

Climatic warming has strongly shifted phenophases in the Northern Hemisphere in recent decades^1,10-12^. Rising temperatures have lengthened the annual growth cycle by advancing leaf unfolding in spring and delaying leaf fall in autumn^13^, albeit with variations among species^14^ and regions^15^. Recent evidence, though, suggests that the sensitivity of spring phenology to warming is decreasing in northern forests^16^ and that the rate of change in plant productivity does not match that of air temperature^17^. Indeed, plant phenology may be acclimated to long-term biogeographical constraints^18-20^ and may be co-limited by several other factors, such as light^21^, water^9,22,23^ and nutrients^24^. These observations suggests that warming does not have the same effect everywhere^25^, which has increased interest in other environmental drivers in recent decades, especially illustrated by multiple debates about the specific role of light (and photoperiodism) in spring phenology^21,26-33^.

## The controversial effect of light: different definitions

How light affects spring phenology remains an open question. Most commonly, its effect is considered via photoperiod, often referred to as daylength. The daylength hypothesis implies that the quality and/or quantity of light is somehow directly sensed by plants through biochemical mechanisms. Some recent studies suggest that the spectral composition of light can indeed influence foliar phenology^34,35^. Light also plays a key role in regulating phytohormones, but the underlying mechanisms remain unknown^32^ and clearly require more investigation. More sporadically, the effect of light has been treated as the sum of insolation over a specific period^36^, for which plants need a specific quotum for a phenological event to occur. The quantity and quality of light depend on plant location, which is the main reason why a response to daylength has often been proposed as a safety mechanism against frost at high latitudes and elevations. Only 35% of the woody species in the Northern Hemisphere, however, depend on daylength as a direct signal for leaf-out^21^, and these species are mainly at mid-to low latitudes

Light effect on spring phenology is still being debated. Recent studies nonetheless suggest a complex interaction between temperature and light. Daytime and nighttime temperatures during winter and spring have an asymmetrical effect on leaf unfolding^37-41^, with a greater weight of temperature during the day^38,42,43^. Whether or not plants are able to sense light, radiation has a physical impact on plants: it affects the temperatures of their tissues. We will use the example of budburst in the following arguments to illustrate that omitting this radiation effect introduces large biases into the interpretation of spring phenological responses based on air temperature.

## The forgotten effects of radiation and wind

Bud temperature (T_bud_) depends on its energy balance^44^. During the day, plant tissues absorb both shortwave (SW, visible and near-infrared) and longwave (LW, infrared) radiation from the sky but also radiation emitted and reflected by the surrounding environment (vegetation, soil) (Figure 1a). Only a fraction (α, absorptivity) of SW radiation will be absorbed depending on bud traits such as color, coating, shape and size (Figure 1b), while most LW radiation will be absorbed by buds. According to the Stefan-Boltzmann law, buds lose energy via LW radiation emission, while they absorb LW radiation emitted from surrounding objects. Finally, a part of their energy is lost by conduction and mostly by convection^45,46^ (e.g. due to wind) while leaves lose an important part of their energy via transpiration.

**Figure 1.**
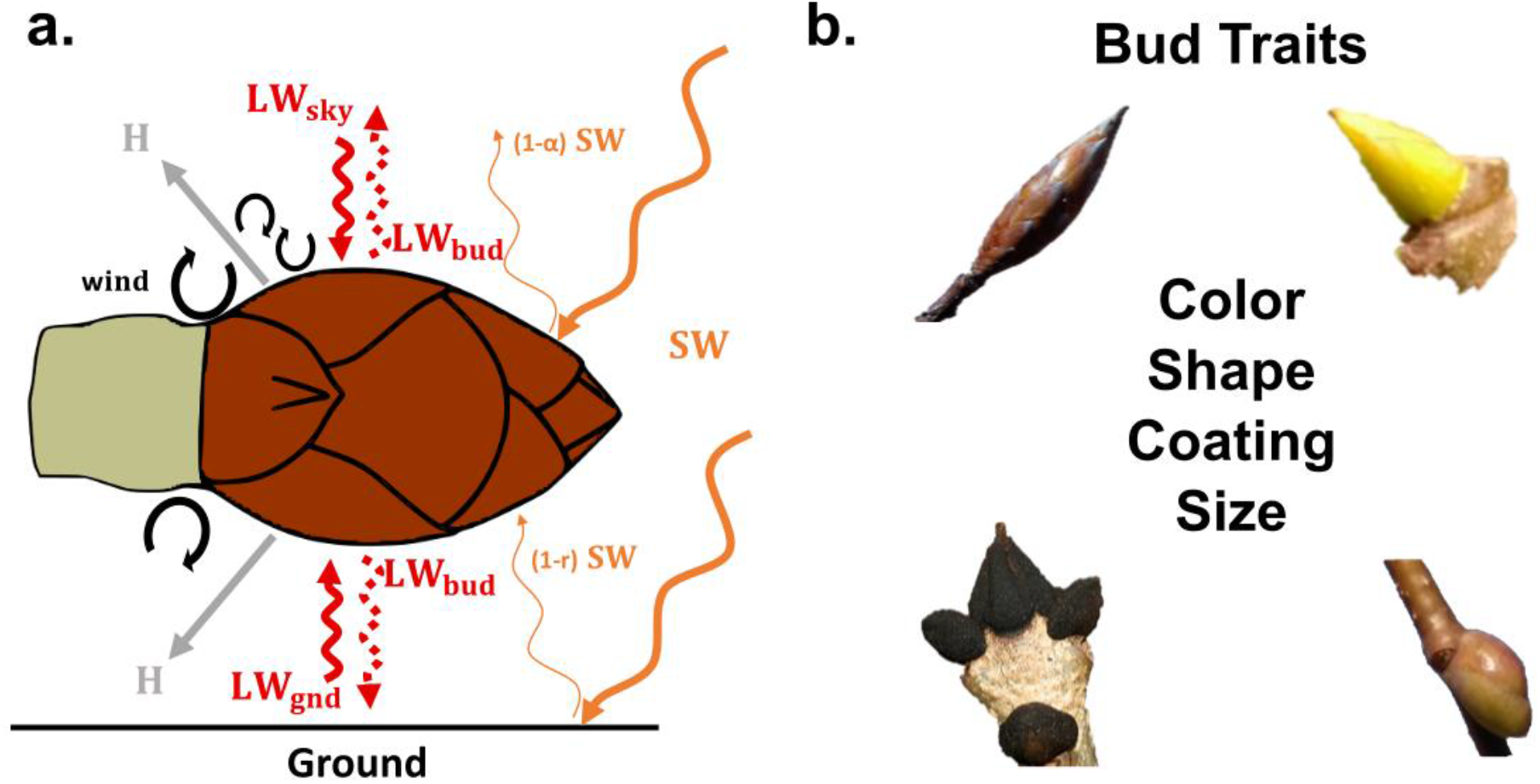
Energy budget of buds and bud traits. **a.** Buds lose energy through convection and conduction (H). Buds absorb incoming shortwave (visible and near-infrared, SW) and longwave (infrared, LW) radiation from the sky (LW_sky_) and the surrounding environment (here simplified as LW radiation from the ground, LW_gnd_). Buds emit LW radiation as a function of their temperature (LW_bud_). Only a fraction (α) of SW radiation is absorbed by buds, depending on the properties of their surfaces. Buds also absorb a small fraction of SW reflected from the ground (1-r). **b.** Illustration of bud traits influencing solar absorptivity, heat conduction and convection processes, and hence, bud temperature.

T_bud_ increases when energy gains exceed losses (Figure 2a) and vice-versa. T_bud_ can thus be lower than air temperatures (T_air_) on clear nights^45^ or because of wind. On the other hand, T_bud_ can be significantly higher than T_air_ during the day. The link between T_bud_ and energy balance has been known for more than 30 years ^44,45^. Since then, all major studies linking temperature and photoperiod to phenological changes, however, have not accounted for the true temperature of plant organs.

**Figure 2.**
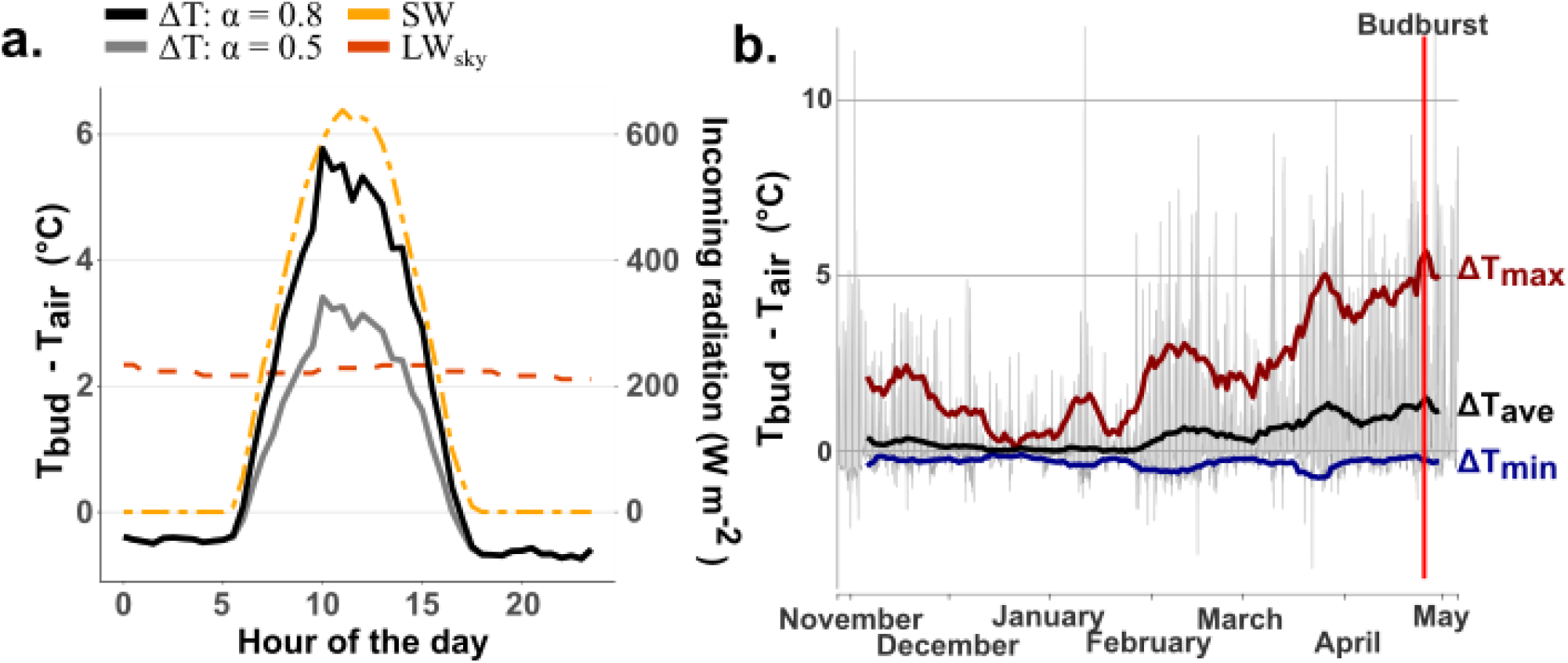
Simulated differences in temperature between buds and the air (ΔT) from energy balance. **a.** Daily variation in ΔT for an exposed bud and a typical day in April for two solar absorptivities: α=0.5 (typical for broadleaves^47^) and α=0.8 (typical for needle leaves^47^). **b** Example of ΔT simulated under idealized conditions (e.g. exposed buds in a deciduous canopy) using meteorological observations for winter and spring collected at the Hesse FLUXNET site^48^ (Beech forest, France). The grey lines represent half-hourly differences in temperature simulated using an energy-budget model. The blue, black and red curves represent the 10-d rolling mean of the minimal (ΔT_min_), average (ΔT_ave_) and maximal (ΔT_max_) temperature differences, respectively. Approximative leaf flushing date is illustrated by the red vertical line (April 25^th^).

## A complex and non-linear response of plant tissue temperature to radiation and wind

What can we expect if we account for the energy budgets of buds in phenological studies? Unfortunately, the lack of *in situ* observations for bud temperature does not allow to answer this question. As part of the reflection, we thus applied existing energy balance approaches^44,45,49^ to explore the potential variability in temperature of an isolated bud (Supplementary material). This situation is well representative of the conditions encountered by sun-exposed buds of a tree and especially of deciduous species (i.e. with no or minimum shading). As a first example, we looked at the variability in T_bud_ estimated from its energy balance and site meteorological observations for an European Beech forest^48^. On average, T_bud_ is expected to be higher than T_air_ during the preseason (~1°C in our example; Figure 2b). Day and night T_bud_ are higher or lower than T_air_ by several degrees. The temperature of buds thus strongly depends on the diurnal radiative cycle and the spectral composition of the light (SW/LW radiation), echoing the observed asymmetrical effect of diurnal temperatures on leaf unfolding^38,42,43^. Applied on four other sites, this approach leads to similar results despite differences in T_bud_ profiles induced by differences in radiation along a latitudinal gradient (Supplementary Figure 1). Spring phenology does not only respond to average preseason temperature, but mainly to the accumulated effect of temperature and its dynamics. It is often assumed that chilling and forcing temperature required for budburst are only effective over specific windows, generally between 0 and 5 °C and over 5°C, respectively. Daily bud temperature variability might thus be the most important factor influencing leaf unfolding, not necessarily its average temperature. We could expect that the difference in extremum temperature sensed by buds over the preceding months (ΔT_min_ and ΔT_max_, Figure 2b) will inevitably affect the apparent forcing and chilling requirement for leaf unfolding.

Accounting for the energy budget of buds for six common species across Europe (Supplementary Figure 2) we also expect a stronger interannual variability in T_bud_ than T_air_, as well as different temporal evolutions over the last decades (Figure 3). In our example, buds are expected to warm faster or slower than air depending on location and species, with 20% and 7% of the sites exhibiting an increase and a decrease in ΔT over 1990-2015, respectively. Even if these trends represent idealized sun-exposed conditions here, we observe that the heterogeneity in ΔT evolution results from a complex and non-linear response to the amount of absorbed radiation and convection processes (Supplementary Figure 3). Because leaf unfolding is earlier in 2015 than in 1990, the average amount of absorbed radiation during the preseason slightly decreased over this period, while most of the interannual variability in ΔT is driven by conduction and convection (i.e. wind). The difference in air-bud temperature and their non-linear and non-proportional relationship suggests that our current interpretation of the apparent bud sensitivity to warming is incorrect.

**Figure 3.**
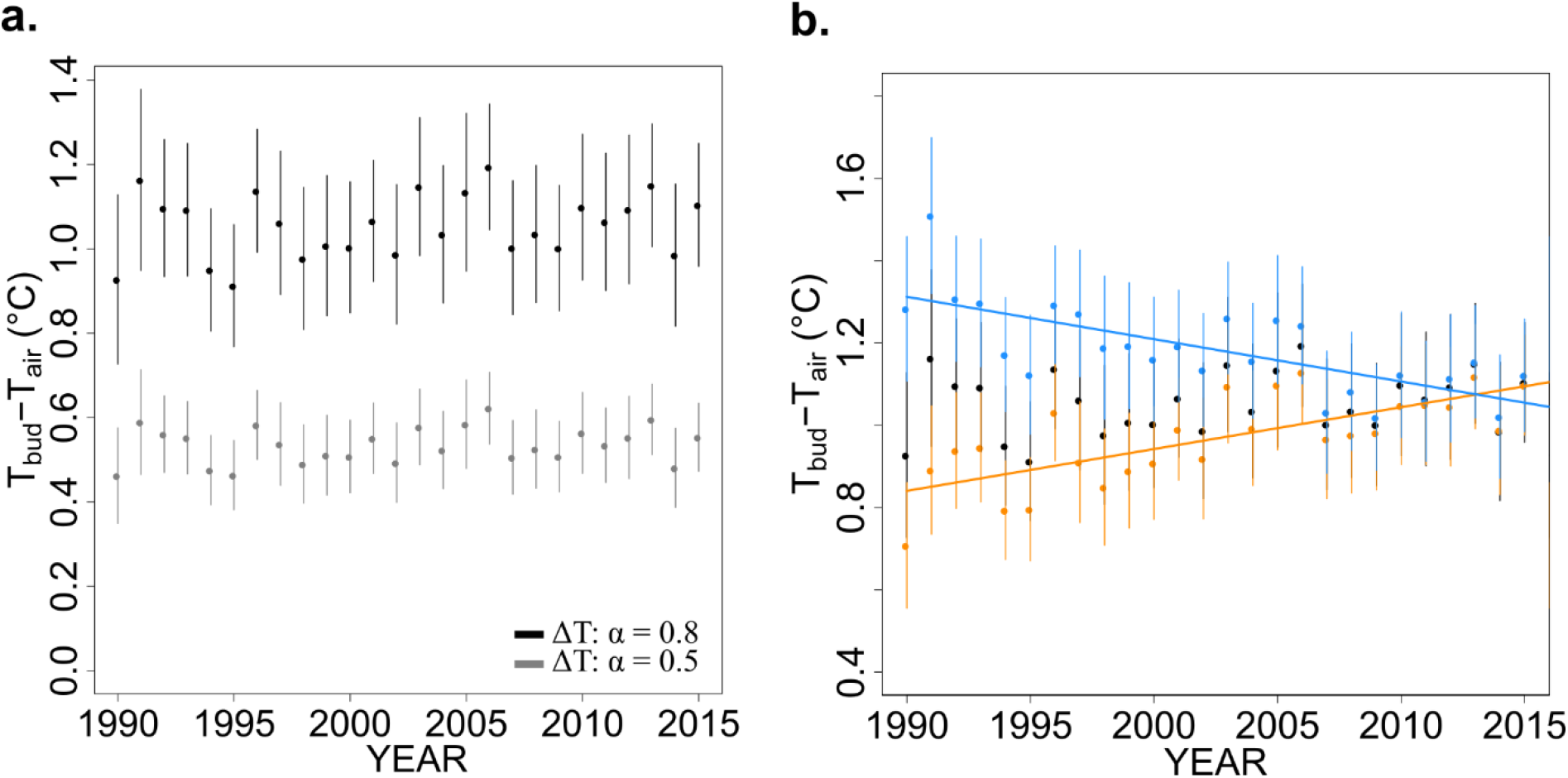
Potential changes in T_bud_-T_air_ over Europe. **a.** Each point corresponds to the mean ΔT simulated for six deciduous species across Europe (1059 sites) under idealized conditions (i.e. sun-exposed buds) using field observation of budburst and global meteorological data (see Supplementary Material). All sites and species were pooled together. Two solar absorptivity values were tested, 0.5 (grey) and 0.8 (black). The error bars represent the spatial and species variability (±1 SD around the mean). **b.** Under these conditions, ΔT is expected to decrease (blue) over the period 1990-2016 for 356 sites*species (7%) and increase for 902 sites*species (orange, 18%) over a total of 5050 sites*species. The black points correspond to all sites pooled together.

## Response to warming will depend on organ properties and local environment

We illustrated the role of bud energy balance through an idealized and constant representation of buds and their environment for all sites and species. Larger spatial and temporal variations are expected due to the effect of topography, ground albedo (e.g. snow, understory), differences in bud traits (Figure 1b) and micrometeorological conditions^50^ that will affect plant tissues energy balance. By affecting the amount of radiation reaching the buds (Figure 1a), varying ground albedo from 0.1 (~wet bare soil) to 0.9 (~snow) leads to a doubling in preseason ΔT (Supplementary Figure 4). Since ground albedo strongly vary in space and over the preseason (e.g snow), we can expect substantial differences in the phenological signal at the regional scale induced by radiation, as already observed from leaf unfolding observations^18^.

Different bud colors or coating will also affect solar absorption of specific wavelengths, while shape and size will modify convection processes and the amount of intercepted radiation (Supplementary Figure 5), and hence, bud temperature. For example, Common Ash (*Fraxinus excelsior*) has black buds while sycamore (*Acer pseudoplatanus*) has green buds and mountain ash (*Sorbus aucuparia*) have dense white trichomes (i.e. hairs) on their surfaces. In our example, a difference in solar absorptivity of 0.3 leads to a doubling in ΔT (Figure 2a, Figure 3a). The differences in bud traits can thus partly account for the observed interspecific differences in heat requirement and apparent sensitivity to temperature. This suggests that the phenological response of plants to warming might be more species-specific than we thought, which should be accounted for in large scale studies.

Despite its central role at the organ level^50,51^, micrometeorology is rarely accounted for in phenology studies because rarely measured, or simply because it is impossible to account for its effect such as in remote sensing analysis or terrestrial biosphere modelling. Instead, phenology studies, either local or regional, often use meteorological and climate dataset with hourly to daily time resolutions. The use of a steady state energy balance is easily justified under such conditions since thermal time constants of tree buds varies between a few seconds to about ten minutes ^52^. Accounting for average preseason radiation and wind conditions might better explain the observed variability in plant phenology than air temperature alone. Here, we only explored spring T_bud_ variability in the case of sun-exposed buds with no shading. Accounting for the potential protecting effect of leaves or needles in evergreen species might substantially attenuate the effects of radiation and wind on intra- and bottom-canopy buds. The concomitant use of high-resolution microclimate data and transient energy budget models will be needed to quantify such effects.

## Towards a better understanding of the environmental control of plant phenology

Drivers of phenological events and light are virtually impossible to separate, because daylength and radiation are strongly correlated with the time of year. Accounting for organ energy balances is thus promising for separating the environmental drivers of phenology using a single approach and potentially for reconciliating the differences observed in the field. Applying existing modelling approaches in the context of sun-exposed buds suggested that air temperature might be an imprecise and biased predictor of bud temperature and more importantly of its variability over the months preceding leaf unfolding, which might introduce biases in the analysis of chilling and forcing requirement for budburst. However, we also showed that bud temperature results from a complex combination of several biotic and abiotic factors, and under certain conditions air temperature might remain a good proxy for bud temperature. The examples we have presented demand the reassessment of past results and interpretations that were solely based on air temperature, however current observations do not allow such reassessment. Bud traits and *in situ* temperature observations are scarcely described in the literature. New experiments and observations are clearly needed for accurately assessing the energy balances of plant organs. Existing studies have mostly focused on leaves, but other organs should also be investigated. Key traits that will need to be measured to assess the interspecific variability of phenology include organ traits influencing solar absorptivity and heat storage, but organ temperatures (i.e. using thermocouples) concomitant with micrometeorological variables will also need to be directly measured. Because buds do not transpire, their energy budget is simpler than for leaves. Properly calibrated, accounting for bud energy balance could improve the accuracy of phenological models that are still unable to predict the spatiotemporal variability of plant dynamics with satisfactory accuracy^53^.

Finally, we stress that energy balance affects the temperature extrema sensed by plants (Figure 2b). The lengthening of the growing season in recent decades has also been associated with an increase in environmental risks. For example, earlier leaf unfolding exposes plants late frost^54-56^ in spring, potentially resulting in dramatic impacts on agriculture^57-59^ and forestry^60,61^. The use of energy balances to study and better predict these environmental risks can provide novel insights into the responses of plants to extreme temperatures and offer more robust predictive tools, which are essential for mitigating the ecological and economic impacts. Temperature of plant organs and their dynamics are still overlooked in both environmental studies and modeling exercises^62^. Energy balance thus plays a key role, not only for plant phenology but also for all other processes since plant tissue temperature will govern key mechanisms such as photosynthesis and respiration and the general functioning of the plant.

## Acknowledgments

The authors would like to acknowledge Dr. Chris Muir and Dr. Renée Marchin Prokopavicius, as well as two anonymous reviewers for their constructive feedbacks on our work. M.P. would like to acknowledge the financial support from the Fonds Wetenschappelijk Onderzoek (FWO; grant no. G018319N) and the H2020 Marie Skłodowska-Curie Actions (LEAF-2-TBM grant no. 891369). J.P. would like to acknowledge the financial support from the European Research Council Synergy grant ERC-SyG-2013-610028 IMBALANCE-P and the Spanish government grant PID2019-110521GB-I00. H.V. acknowledges the support from the European Research Council Starting Grant 637643 TREECLIMBERS.

## Supplementary information

Already existing models and data were used to support this perspective. This work did not attempt to develop, nor validate, a new model. The full description of the energy budget model (and underlying assumptions) as well as supplementary figures and tables are provided in the Supplementary information. The R code of the model of energy budgets and data sets used to generate the figures and analysis of this manuscript are available from Github: https://github.com/mpeaucelle/Tbud

A version of the git repository is archived on Zenodo at https://zenodo.org/record/4173415.

