## Supplementary material for "Warming sensitivity of spring phenology of deciduous species lies in their bud energy budget"

**Detailed description of the bud energy budget model**

We implemented a simplified energy budget model for buds based on Landsberg et al. (1974)[1], Hamer (1985)[2] and equations from Muir (2019)[3]. We refer to Table S1 for parameter values and units.

For an isolated bud, the amount of absorbed incoming radiation ($R_{abs}$) is balanced by the thermal infrared radiation loss ($LW_{bud}$) and the energy lost by conduction and convection, generally called the sensible heat flux ($H$). The thermal time constant of buds varies between a few seconds to a dozen of minutes[4], meaning that bud temperature responds quite rapidly to local changes. Because most phenology studies use meteorological data from local stations or gridded datasets, here we simulate bud temperature by considering that the energy balance is close to equilibrium at a time scale of a few minutes, which is consistent with temporal resolution of meteorological observations from FLUXNET sites (30 min) and CRU-JRA[5] (6 h):

$$R_{abs} = LW_{bud} + H \qquad (1)$$

The amount of absorbed energy by buds is the sum of incoming shortwave (SW, visible and near-infrared) and longwave (LW, infrared) radiations:

$$R_{abs} = \alpha_{sw}(1 + r)SW + \alpha_{lw}(LW_{sky} + LW_{gnd})/2 \qquad (2)$$

where $\alpha_{sw}$ is the bud absorptivity to SW; $r$ is the fraction of SW reflected by the ground and $\alpha_{lw}$ is the bud absorptivity to LW, which is here defined as the average of LW coming from the atmosphere $LW_{sky}$ and coming from the ground $LW_{gnd}$.

LW emitted by surrounding objects such as branches were considered equal to LW emitted from the ground and the sky. We set $\alpha_{lw}$ to 0.97, which corresponds to the average absorptivity for wood and leaves [6–8]. Since no data were available in the literature, we tested two different values of $\alpha_{sw}$, 0.5 and 0.8, corresponding to values commonly used for broadleaves and needleleaves, respectively. We set $r$ to 0.2, which corresponds to a reasonable value for the fraction of SW reflected by grasses. Of course, $r$ will strongly vary according to the albedo of the ground (e.g. understory/grass, forest litter, snow, etc.), which was simplified here for our perspective paper.

In our example, we simplified the estimation of $LW_{gnd}$ by considering that ground temperature equals air temperature:

$$LW_{gnd} = \sigma\varepsilon T_{air}^4 \qquad (3)$$

where $\sigma$ is the Stefan-Boltzmann constant and ε is the bud emissivity to longwave radiations (which is equal to $\alpha_{lw}$ since plant material tends to behave like a black body in the IR spectrum[6]). This is an important oversimplification to keep in mind since ground temperature will strongly depend on soil type, vegetation and snow cover, soil humidity, but also the fact that soil heat capacity is generally higher than air.

Buds lose thermal infrared radiation proportionally to their temperature as:

$$LW_{bud} = \sigma \varepsilon T_{bud}^{4} \qquad (4)$$

where $\sigma$ is the Stefan-Boltzmann constant and $\varepsilon$ is bud emissivity to longwave radiations.

Finally, the sensible heat flux depends on the air to bud temperature gradient and is formulated as in Muir (2019)[3]:

$$H = P_a c_p g_b (T_{bud} - T_{air}) \qquad (5)$$

where $P_a$ is the density of dry air; $c_p$ is the specific heat capacity of air at constant pressure and $g_b$ is the boundary-layer conductance to heat.

$$P_a = \frac{2P}{R_{air}(T_{bud} - T_{air})} \qquad (6)$$

where P is the atmospheric pressure, $R_{air}$ is the specific gas constant for dry air.

$$g_b = \frac{D_h Nu}{d} \qquad (7)$$

where Nu is the Nusselt number, Dh is the diffusion coefficient of heat in air and d is the bud diameter.

$$D_h = D_{h,0} \left(\frac{T}{273.15}\right)^{eT} \frac{101.3246}{P} \qquad (8)$$

$D_h$ is function of temperature and pressure, $D_{h,0}$ corresponds to $D_h$ at 0°C and eT is the temperature dependence of diffusion.

The Nusselt number is estimated as a mixed convection such as:

$$Nu^{3.5} = Nu_{forced}^{3.5} + Nu_{free}^{3.5} \qquad (9)$$

with

$$Nu_{forced} = e + aRe^{b} \qquad (10)$$

and

$$Nu_{free} = f + cGr^{d} \quad (11)$$

*Re* and *Gr* are the Reynolds and Grashof numbers, respectively; *a, b, c* and *d* are constants that are dependent of the flow regime.

In our example, we simplified bud's shape as a spherical object with a diameter of 5 mm to computes the boundary-layer conductance to heat. Please refers to Hamer (1985)[2] for a detailed discussion of this assumption, as well as empirical formulations for the convective heat transfer of apple buds. Condition for laminar and turbulent flows, as well as constants *a, b, c* and *d* for a spherical object, as well as for objects of various forms, can be found in Monteith and Unsworth (2013)[9].

In addition, we performed two simulations varying ground albedo from 0.1 (~wet bare soil) to 0.9 (~snow) and bud diameter from 5 to 13 mm (Supplementary Figure 4 and 5)

The bud energy model simulates bud temperature **in idealized conditions and at equilibrium**, without distinction between species and sites and several simplifications as described above.

**Special Note:** a proper calibration of species and site properties (e.g. albedo) is needed to use this model for predictions. Here, we used already existing models and assumptions for the only purpose of exploring the expected variability in bud temperature with environmental conditions and for an isolated object representative of apical buds in a tree. Model development and validation was not intended here and will require *in situ* observations of bud traits and temperature data.

Because available meteorological observations have a time resolution between 30 min and 6h, here we used a steady state approach to simulate bud temperature. Using such modeling approach with high resolution micrometeorological observations will lead to better results with a transient energy model[6] taking into account for the thermal time constant of buds.

**Data availability**

Half-hourly forcing meteorological data (Figure 2, Supplementary Figures 1, 4 and 5) were downloaded from the FLUXNET2015 dataset at https://fluxnet.fluxdata.org/data/fluxnet2015-dataset/. Historical climate data from CRU-JRA[5] at a spatial resolution of 0.25° and at a temporal resolution of 6 h (Supplementary Figures 3 and 4) were downloaded at https://catalogue.ceda.ac.uk/uuid/13f3635174794bb98cf8ac4b0ee8f4ed.

*In situ* leaf unfolding data (Figures 3 and Supplementary Figure 4) for Common alder (*Alnus glutinosa*), horse chestnut (*Aesculus hippocastanum*), silver birch (*Betula pendula*), European beech (*Fagus sylvatica*), European ash (*Fraxinus excelsior*) and pedunculate oak (*Quercus *robur*) were downloaded from the Pan European Phenology network (http://www.pep725.com/). Phenological observations followed the Biologische Bundesanstalt, Bundessortenamt und Chemische Industrie (BBCH) code, with leaf unfolding corresponding to BBCH = 11. Only sites with more than 20 years of observation over the 1990-2015 period were used, corresponding to 5050 sites*species in total, covering 1059 sites (Supplementary Figure 2).

**Table S1 | List of variables and parameters**

| Variable | Description | Units |
|---|---|---|
| Rabs | Amount of absorbed radiation by buds | W $m^{-2}$ |
| SW | Shortwave radiation (visible and near infrared) | W $m^{-2}$ |
| LW | Longwave radiation (infrared) | W $m^{-2}$ |
| H | Sensible heat flux | W m-$^{2}$ |
| $\alpha_{sw}$, $\alpha_{lw}$ | Absorptivity to SW and LW (0-1). | unitless |
| ε | Emissivity to LW (0-1) | unitless |
| r | Fraction of reflected SW by the ground (0-1) | unitless |
| T | Temperature | K |
| P | Atmospheric pressure | kPa |
| $P_a$ | Density of dry air; | g $m^{-3}$ |
| $g_b$ | Boundary-layer conductance to heat | m $s^{-1}$ |
| $D_h$ | Diffusion coefficient for heat in air | m² $s^{-1}$ |
| Re | Reynolds number | unitless |
| Gr | Grashof number | unitless |
| Nu | Nusselt number | unitless |
| **Constant parameters** | | |
| σ | Stefan-Boltzmann constant | 5.67 $10^{-8}$ W $m^{-2}$ $K^{-4}$ |
| $c_p$ | Specific heat capacity of air at constant pressure | 1.01 J $g^{-1}$ $K^{-1}$ |
| $R_{air}$ | Specific gas constant for dry air | 287.058 J $kg^{-1}$ $K^{-1}$ |
| $D_{h,0}$ | Diffusion coefficient for heat in air at 0°C | 19.0 10-6 m² $s^{-1}$ |
| eT | Exponent for temperature dependence of diffusion | 1.75 |
| a | Constant to estimate Nu for a smooth sphere[9] | If Re > 50; a = 0.34<br>If Re > 300; a = 0.54 |
| b | Constant to estimate Nu for a smooth sphere[9] | If Re > 50; b = 0.6<br>If Re > 300; b = 0.5 |
| c | Constant to estimate Nu for a smooth sphere[9] | For $Gr^{0.25}$ < 220; c = 0.54 |
| d | Constant to estimate Nu for a smooth sphere[9] | For $Gr^{0.25}$ < 220; d = 0.25 |
| e | Constant to estimate Nu for a smooth sphere[9] | If Re > 50; e = 0<br>If Re > 300; e = 2 |
| f | Constant to estimate Nu for a smooth sphere[9] | For $Gr^{0.25}$ < 220; f = 2 |

### Supplementary Figures

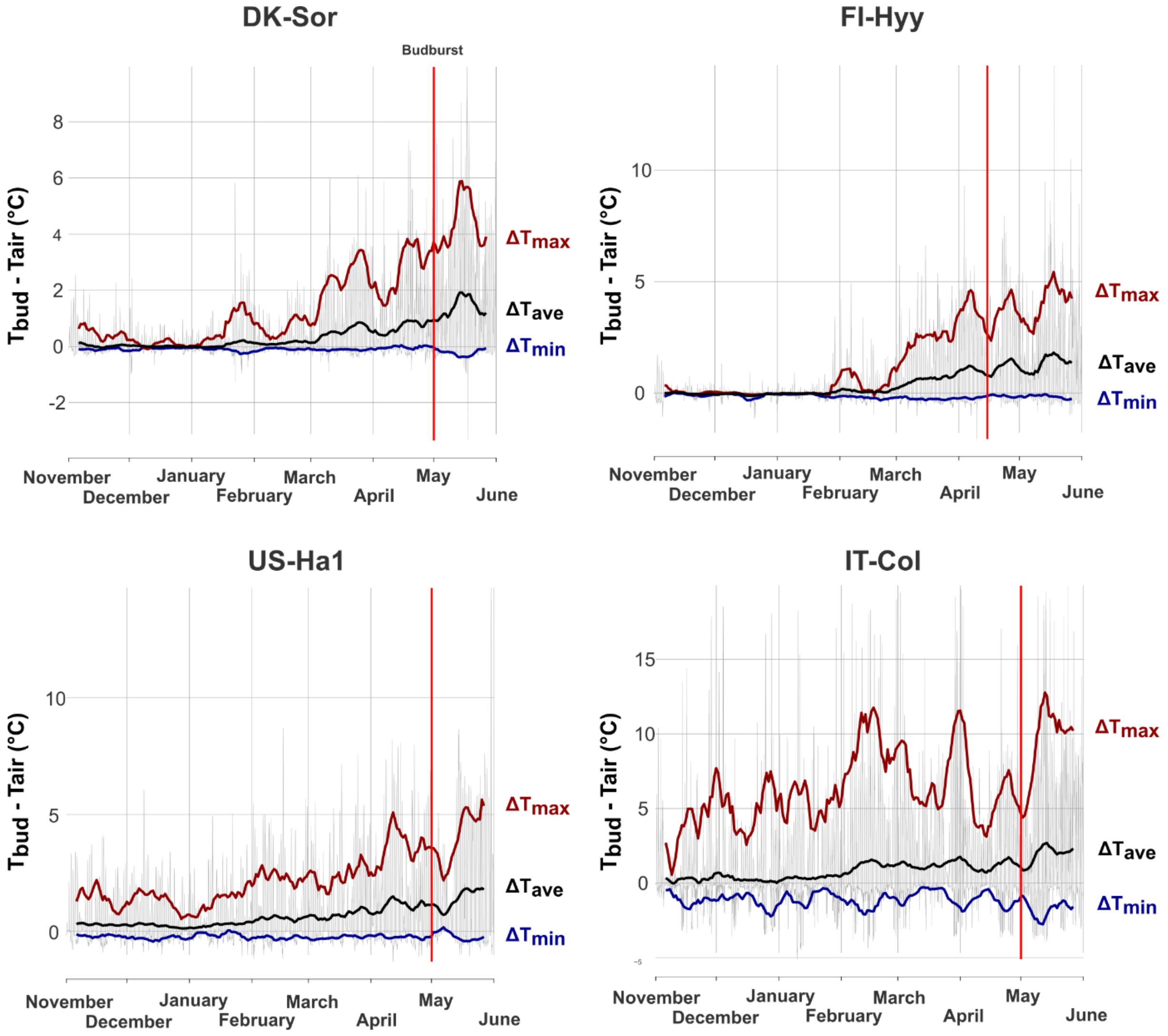


**Supplementary Figure 1 | Spatial variability in bud temperature.** Temperature difference (ΔT in °C) between buds and air simulated for four different FLUXNET sites during the year 1998: DK-Sor (Deciduous Broadleaf Forest; 55.48 °N, 11.64 °E), FI-Hyy (Evergreen Needleleaf forest; 61.85 °N, 29.3 °E), US-Ha1 (DBF; 42.53 °N, 72.17 °W) and IT-Col (DBF; 41.85°N, 13.59°E). The grey lines represent half-hourly differences in temperature simulated using the energy-budget model. The blue, black and red curves represent the 10-d rolling mean of the minimal ($\Delta T_{min}$), average ($\Delta T_{ave}$) and maximal ($\Delta T_{max}$) temperature differences, respectively. Approximative budburst is represented by the red vertical line. Note that these simulations represent idealized conditions and use the same parameterization without distinction between sites and species.

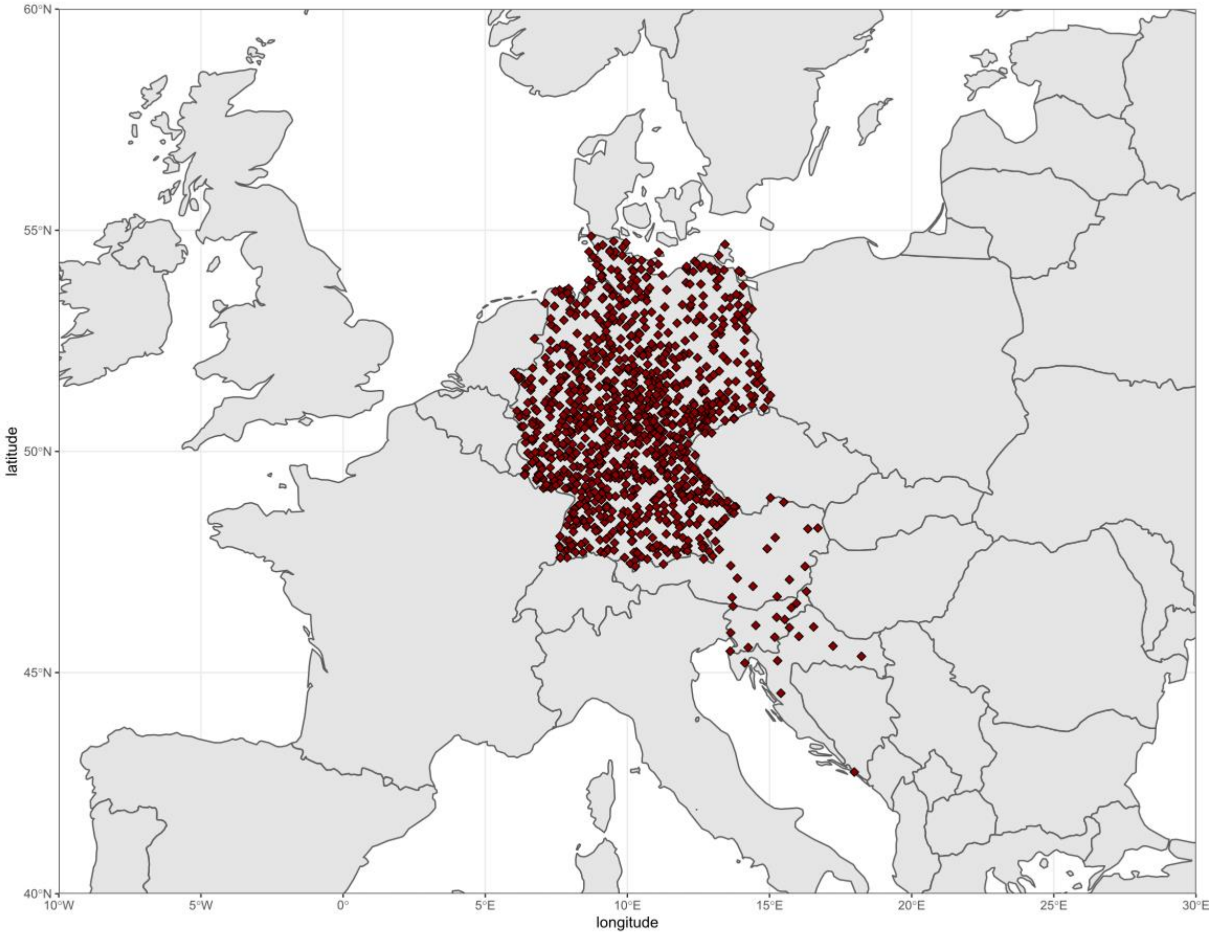


**Supplementary Figure 2 | Distribution of sites with more than 20 years of leaf unfolding records over 1990-2015.** Sites including data for Common alder (*Alnus glutinosa*), horse chestnut (*Aesculus hippocastanum*), silver birch (*Betula pendula*), European beech (*Fagus sylvatica*), European ash (*Fraxinus excelsior*) and pedunculate oak (*Quercus robur*) for the PEP database.

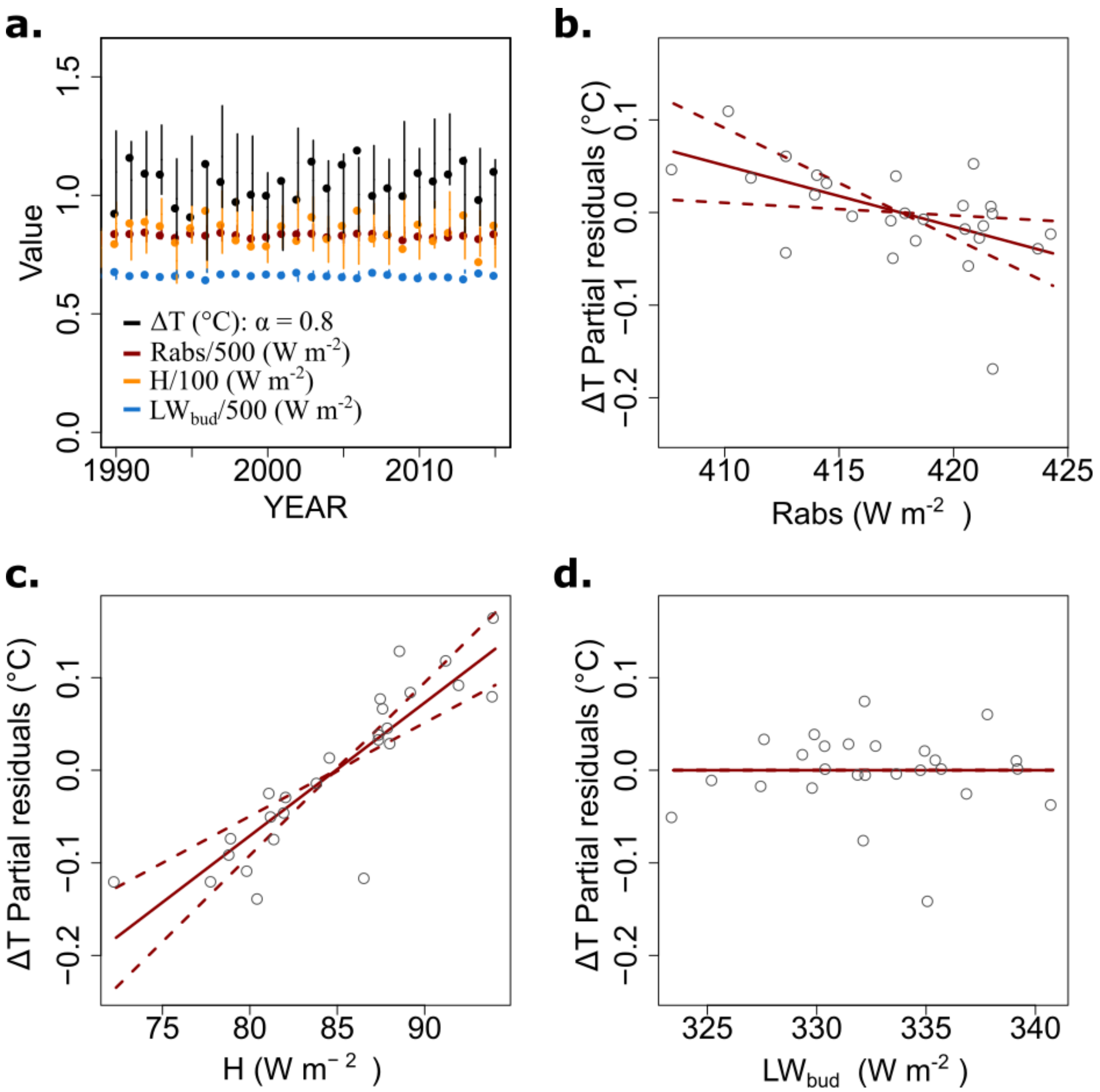


**Supplementary Figure 3 | Weight of each component of the bud energy budget in simulating bud temperature. a.** This figure illustrates that the response of the bud-air temperature difference (ΔT) is a complex nonlinear response to Rabs (red), H (orange) and $LW_{bud}$ (Blue). For more clarity, energy components were divided by 500, 100 and 500 for Rabs, H and $LW_{bud}$, respectively. Each point corresponds to the mean ΔT simulated for six species across Europe (1059 sites) under idealized conditions using field observation of budburst and global meteorological data. All sites and species were pooled together. Solar absorptivity to shortwave radiation is set to 0.8. The error bars represent the spatial and species variability (±1 SD around the mean). Panels **b**, **c** and **d** represent the weight of Rabs, H and $LW_{bud}$ in explaining ΔT variability, by showing the relationships of their respective partial residuals in the multiple linear regression $\Delta T = Rabs + H + LW_{bud}$ ($R^2 = 0.85$) using global yearly averages. We observe that H and Rabs account for most of the variability in ΔT. Red full lines represent the slope coefficient in the linear regression. Dashed lines represent the standard error associated to each coefficient.

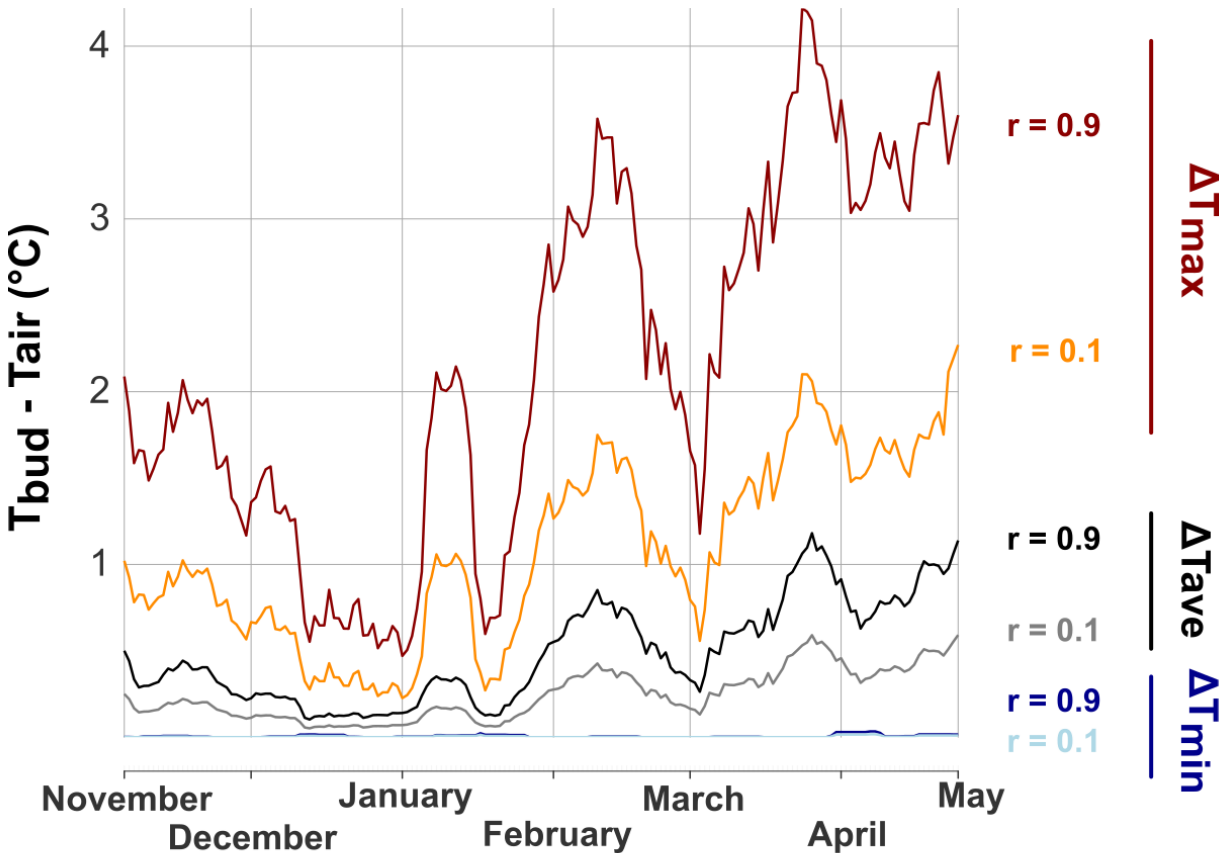


**Supplementary Figure 4 | Impact of albedo on simulated bud temperature.** Temperature difference (ΔT in °C) between buds and air simulated for two different ground albedo, r = 0.1 (~wet bare soil) and r = 0.9 (~snow). Temperature difference was simulated using meteorological observations for winter and spring collected at the Hesse FLUXNET site (Beech forest, France; see Figure 2). The differences for maximum (red), average (black) and minimum (blue) temperatures are illustrated. Here, bud solar absorptivity to shortwave radiation and bud diameter were set to 0.8 and 5 mm, respectively.

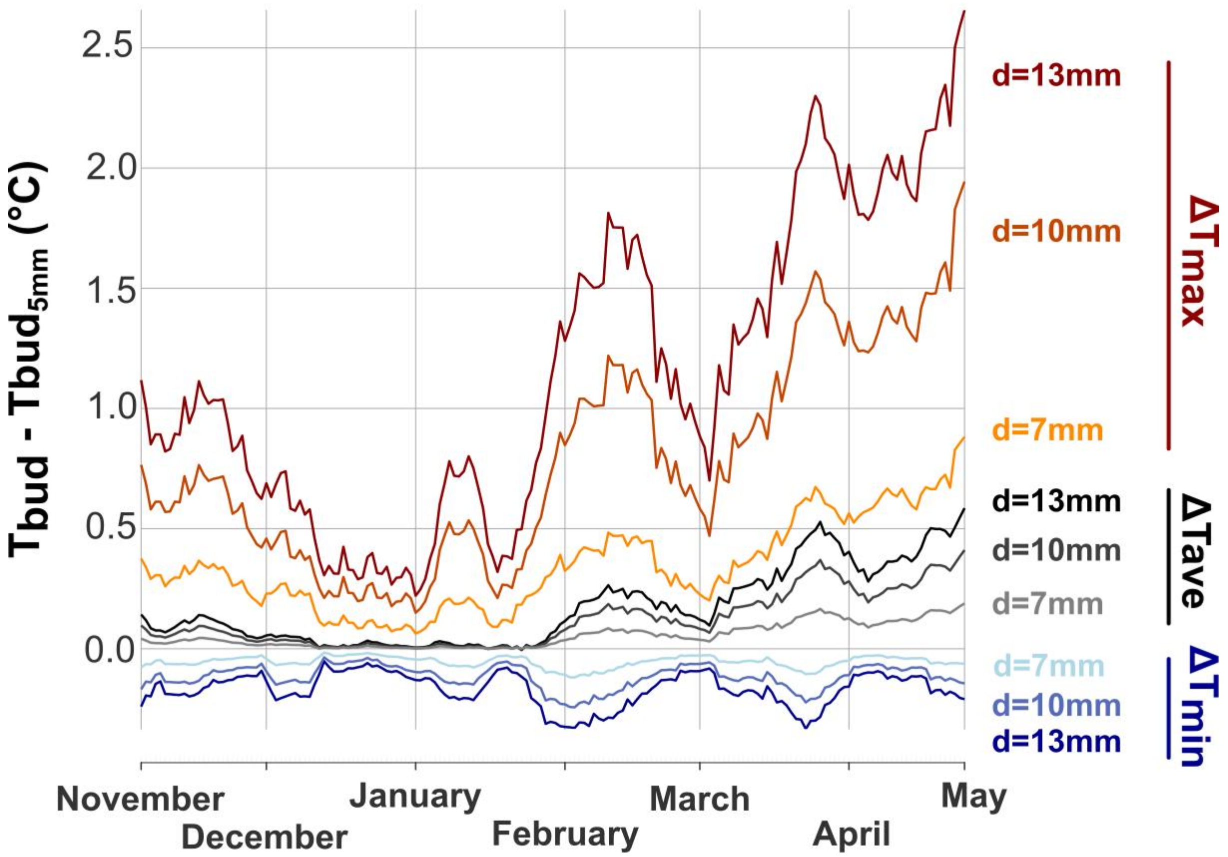


**Supplementary Figure 5 | Impact of bud size on simulated bud temperature.** Temperature difference (ΔT in °C) between buds with a diameter (d) of 7, 10 and 13mm and buds with a diameter of 5mm (default value used in the manuscript). Temperatures were simulated using meteorological observations for winter and spring collected at the Hesse FLUXNET site (Beech forest, France; see Figure 2). The differences for maximum (red), average (black) and minimum (blue) temperatures are illustrated. Big buds exhibit larger temperature variations than small buds. Here, bud solar absorptivity to shortwave radiation and ground albedo were set to 0.8 and 0.2, respectively. Please note that these results represent the average change in bud temperature with a steady state model. With a low resolution of 30 min in meteorological forcing compared to the relatively small thermal time constant of buds[4], big buds have the time to accumulate more energy than small buds. Using a transient model, calibrated with site data and high-resolution micrometeorological observations accounting for the sudden changes in wind and radiation might lead to different results.
